# Controlling water hyacinth infestation in Lake Tana using Fungal pathogen from Laboratory level upto pilot scale

**DOI:** 10.1101/2020.01.14.901140

**Authors:** Adugnaw Admas, Samuel Sahile, Aklilu Agidie, Hailu Menale, Tadelo Gedefaw, Menberu Teshome

## Abstract

Water hyacinth (Eichhornia crassipes) is one of the most dangerous aquatic weeds for Lake Tana and other water body in Ethiopia. To reduce its invasion biological, chemical and physical control methods can be used. Use of natural biological enemies of the weed to discourage its propagation is one of the best recommended options by scientfic society. Among them, there are more fungi naturally a pathogen for water hyacinth and other plants. To use those patogenes to manage water hyacinth infestation in Lake Tana infected plant material by fungi were collected from three weredas (Amba Gyorgese, Dabat and Debarke) around Gondar at 20 Peasant associations (PAs) since Novmber 2015. The collection was done from infected Faba bean leaves and roots. All isolated fungus was attempted to infect the collected healthy water hyacinth in laboratory and green house. Among isolated fungus species Rhizoctonia solani, Aspergillus flatus, Tricothcium roseum, Fusarium spp and Aspergillus niger fungi show high moderate disease severity on the healthy water hyacinth at temporarey green house and laboratory. Disease severity scale was recorded using modified NAHEMA et al. (200). By following those experiments to show its efficiency, the effective pathogens on laboratory and green house were released to 16 m2 open ponds since September 2016, in University of Gondar. In this study, we have recorded scientific data that shows the fungi were high potencial to attack healthy water hyacinth at above 26 oc and at less than 25 % humidity. From this research also we have observed the most infected water hyacinth by fungi have not produced flower and it can not re generate by seed in the next propagation sesaon.Finally, before directly release the fungi on Lake Tana its impacts were studied in the Goregora, at Kuame Michel kebela for a year in open ponds and in controlled wet land areas that not linked to the Lake by taking some common aquatic plants and fish from the Lake. Fortunatelly, those fungi have not impact on aquatic plant like Echinochloa and Cyperus papyrus grass, water quality and fish.

## 1. INTRODUCTION

Lake Tana is occurred in the highlands of north-western Ethiopia.It is the country’s largest freshwater body and the third largest lake in the Nile Basin. Also, it is the source of the Blue Nile, and its basin is one of the most important catchments in the Nile Basin (Bijan and Shields, 2011). The LakeTana basin has high significance to the economy and politics of Ethiopia. It also greatly influences the livelihoods of millions of people in the lower Nile Basin. Historically, there was a large area of Afromontane forest and many indigenous plant species in the Lake Tana basin; 172 woody species were observed in the basin, many of which were indigenous species (IFAD, 2007). There are also large areas of wetlands and seasonally flooded plains, which provide multiple services to the local community and serve as a home for many endemic bird species (Bijan and Shields, 2011). Lake Tana has a mean depth of 9 metre and maximum depth of 14 meter, length 78 kms and width of 68kms (Amare,2011). The basin (Lake Tana Basin and Nile Valley) was also the repositories of ancient indigenous culture, linguistics, history and ancient civilization. Some thousand years ago, science and technology in Medicine, Mathematics, Agriculture, Architecture, communication or writing system, commerce, etc.) were practiced in the Nile basin and it is assumed that the current world’s technology is originated from the Nile basin and African rift valley (UNEP, 2006; Wikipedia). The lake also balances the climatic condition of the region. The northern pick (i.e. Rase Dejene) and Lake Tana is the green belt which prevents the spread of the Sahara desert and offer wetter climates. The climate of the Eastern Afromontane Hotspot has been relatively constant over recent geological history due to high levels of biodiversity, endemism and thermo — coolant factors of the lakes and mountains (Balm ford et al., 2001; Borgess et al., 2007). In addition, the Lake supports hydro electric powers. In this regard, the lake plays a role to control environmental pollution. Hydropowers are most importantly clean energy sources with relatively negligible production of noxious gases or solid/ liquid wastes, and therefore the dam can largely prevent environmental pollution, climate change, global warming, and related public health consequences. Levels of greenhouse gases (GHG) emission from hydro-power are relatively low (Kaunda et al., 2012). The life cycle GHG emission factors for hydro-power technologies are around 15–25 g CO2 equivalent per kWhel. These are very much less than those of fossil-fuel power generation technologies which typically range between 600–1200 gram CO2 equivalents per kWhel (Lenzen, 2008). As a Portuguese missioner Manoel de Al media in the early 17 centuries wrote in the center of Lake Tana 21 islands were exsit, among them seven to eight of which had monasteries. In most monstery,Ethiopian emperors and treasures of the Ethiopian Churchs were kept among them Dega Estifanose, Ura Kidane mihrit, Narga Selassie, Keberan Gabriel, Medhanelm of Rema, kota mar yam and Mertola maryam were found, also when the emperors become died their tombs are kept in those isolated monasteries among them Yekuno AMLA, Davit I, ZaraYaqob, Za Den gel, and Fasilide’s tombs are kept (Beck ham etal,1954) also, Ethiopian Orthodox religious followers believe on the islands of Tana Qirkose the Virgin Maryam had rest from her journey when she back from Egypt and Christianity to Ethiopia is raise in Tana Cherqose (Paul B. Hence,2000). Also,there is a believe by society those lives in the edges of the lake when Virgin Mariam returne from Tana Qirkose she was take rest in two place,those are Mahdere Mariam in South Gondar zone,Deberetabor district and at Mariam chuarche in Alem Saga forest, Fogera Districts. Lake Tana is not only the economic and cultural source of Ethiopians but also it is the economic source of Egypt and Sudan, but now Water hyacinth weed is the challenges of this lake. Since 2011 it is observed in edges of the Lake and in 2014 it spreads on more than 50,000 hectors of the water surfaces from total area of 360, 000 hectares (Wassie Anteneh et al., 2015) but from 2015-2019 till this study clear data is not in a hand that shows the amounts of the weed infestation in Lake Tana. Water hyacinth is a free — floating perennial plant native to tropical and sub-tropical South America. It rises above the surface of the water as much as 1 meter in height and have 80 cm root below the surface of water. The leaves are 10–20 cm wide and float above the water surface. They have long, spongy and bulbous stalks. It reproduces primarily by way of runners or stolons. Each plant additionally can produce thousands of seeds each year and seeds can remain viable for more than 28 years (Sullivan et al., 2012). It also doubles their population in two weeks. International Union for Conservation of Nature(IUCN’s) has listed this species as one of the 100 most dangerous invasive species (Teller et al., 2008) and the top 10 worst weeds in the world (Shanab et al., 2010). The water hyacinth appeared in Ethiopia in 1965 at the Koka Reservoir and in the Awash River(Rezene Fessehaye, 2005). It affects navigation, water flow, recreational use of aquatic systems, and causes mechanical damage to hydroelectric systems (Navarro and Phiri, 2000). It is also responsible for drastic changes in the plant and animal communities of fresh water environments and acts as an agent for the spread of serious diseases in tropical countries. The impact of Eichhornia crassipes on the physico–chemical characteristics of the water in general are declines in temperature, pH, biological oxygen demand (organic load), and nutrient levels (Rai and Munshi 1979). Biological control using plant pathogens has been found to be highly effective against water hyacinth under experimental conditions (Shabana, 1997). Several highly virulent fungal parasites are known to cause diseases of water hyacinth (Charlatan, 1990). Among the known pathogens are Alternaria eichhorniae, PERCO sporarodmanii and Fusarium chlamydosporum have been studied to a significant extent (Charlatan, 1990; ANENA et al., 1993). Hence, the aim of this study were to investigate the bio control potential of fungus on water hyacinth and assess its impacts on common biodiversity of Lake Tana.

## 2. MATERIAL AND METHODS

### 2.1. Study Area

The study areas were located in northern part of Lake Tana in Dembiya woreda, Gorgora at 1190SE12015’45’’ N37018’11’’E

### 2.2. Sampling method

Surveying Diseased Faba bean leaves (showing browning, wilting, yellowing, spots, blights, or combinations and its roots were collected randomly from 3 districts at 20 kebeles using plastic bags (Ambagyrorgise, Dabat and Debarke around Gondar and healthy water hyacinth was collected in Lake Tana for treatment. The fungus was isolated using potato dextrose agar medium from infected leaf and root. Fungal pathogens are able to infect various plant parts such as roots, stems, leaves, flowers and fruits, inducing characteristic visible symptoms like spots, blights, anthracite and wilts. Collected infected parts of Faba bean was cut into small pieces. After washing the tissues thoroughly in sterile water, the causal fungi are isolated from plant tissues exhibiting clear symptoms. The infected tissues along with adjacent small unaffected tissue are cut into small pieces (2–5 mm squares) and by using flame-sterilized forceps, they are transferred to sterile petridishes containing 97% ethanol used for surface sterilization of plant tissues. The plant parts were transferred to PDA plates and incubated for 5-7 d for the complete growth of fungi. The fungi were identified according to cultural characters described by Gilman (1957), Barnett and Hunter (1972) and Nelson et al. (1982).

## 3. Water hyacinth plant pathogncity test in temporary shade

Healthy water hyacinth plants were collected from natural infestations of Lake Tana and maintained in a sterilized condition. Water hyacinth plants were kept in plastic pot filled with water and moisture containing sandy soil. In 250 ml Erlenmeyer flask, each contains 100 ml malt extract broth (MEB). were sterilized at 121 0_C_ for 20 minutes and inoculated after cooling 7 - 10 day old cultures of isolated fungi. The inoculums spore suspension was incubated at 25 0_C_ on rotary shaker for 5 - 7 days. The resulted 10 ml mycelium suspension were diluted by 20 ml of distiled water for each replicate, for three replicate 30 ml fungal suspension diluted with 60 ml of distiled water. For comparison of pathogen city, each suspension was then liberally applied to the surface of water hyacinth plants using a hand sprayer and all treated water hyacinth plant was covered by polythene tube sheet up to disease symptoms observed on leaf to minimize the disspercity across each other treated plants. Control plants were placed under the same conditions but without addition of the antagonists. Data collection commenced immediately disease symptoms appeared and continued weekly up to a five-week period. Plants with disease symptoms were recorded and scored to give disease incidence and severity. The disease incidence (DI) was taken as the percentage number of leaves on the plant that exhibited disease symptoms. This was measured by observing all the leaves of the inoculated plant in each pot and calculated as percentage of the total number of leaves on the plant. Disease severity (DS) was determined for each leaf on a scale of 0 to 9, where 0 = healthy, and 9 = 100 % diseased (Freeman and Charlatan, 1984). Values for individual leaves were summed and averaged to derive DS for a whole plant. Finally, isolates were categorized into five groups: “N,” isolates that did not cause any significant damage or infection, “Mild,” isolates that caused less than 25 % damage to the leaf area; “Low Moderate” isolates caused 26-50 % damage to the leaf area; “ High Moderate,” isolates that damaged 51-75 % of the leaf area and “Severe,” fungi that cause greater than 75 % damage to the leaf area. 3.4. Data analysis Data was carried out using Statistical Package for Social Science (SPSS) version 16.0. Descriptive Statistics were employed to present the percentage of isolates. All the presently performed analyses were carried out in duplicate and the correlation were calculated. Mean separation were carried out using the least significant differences (LSD) between means for the different treatments. P = 0.05 was taken as statistically significant association.

### 3.1. Water hyacinth pathogencity test by fungi in pond

The pathognecity of the fungi on water hyacinth were assese at 16 m2 open ponds by cultivating healthy water hyacinth.The fungus that showed high pathogen cities in green house experiment were Rhizoctonia solani., Aspergillus flatus, Tricothcium rose um, Aspergillus niger grown in 250 ml flasks each contain 100 ml of PDA broth. By using pressure sprayer the fungi released on prepared ponds in the areas of 16 metre square. The data were collected by divided four quadrants and regularly count the infected leaf in each month (September 2016 – April 2017).

## 4. Results and discussion

Out of 230 isolating fungi from three districts, seven fungi species were affected the healthy water hyacinth in green house experiment. Those fungi were *Tricothecium roseum*, *Aspergillus Flaves, Trichoderma spp1, Fusarium spp, Rhizocotonia spp, Aspergillus niger* and *Trichdoerma spp*.

**Table 1:**
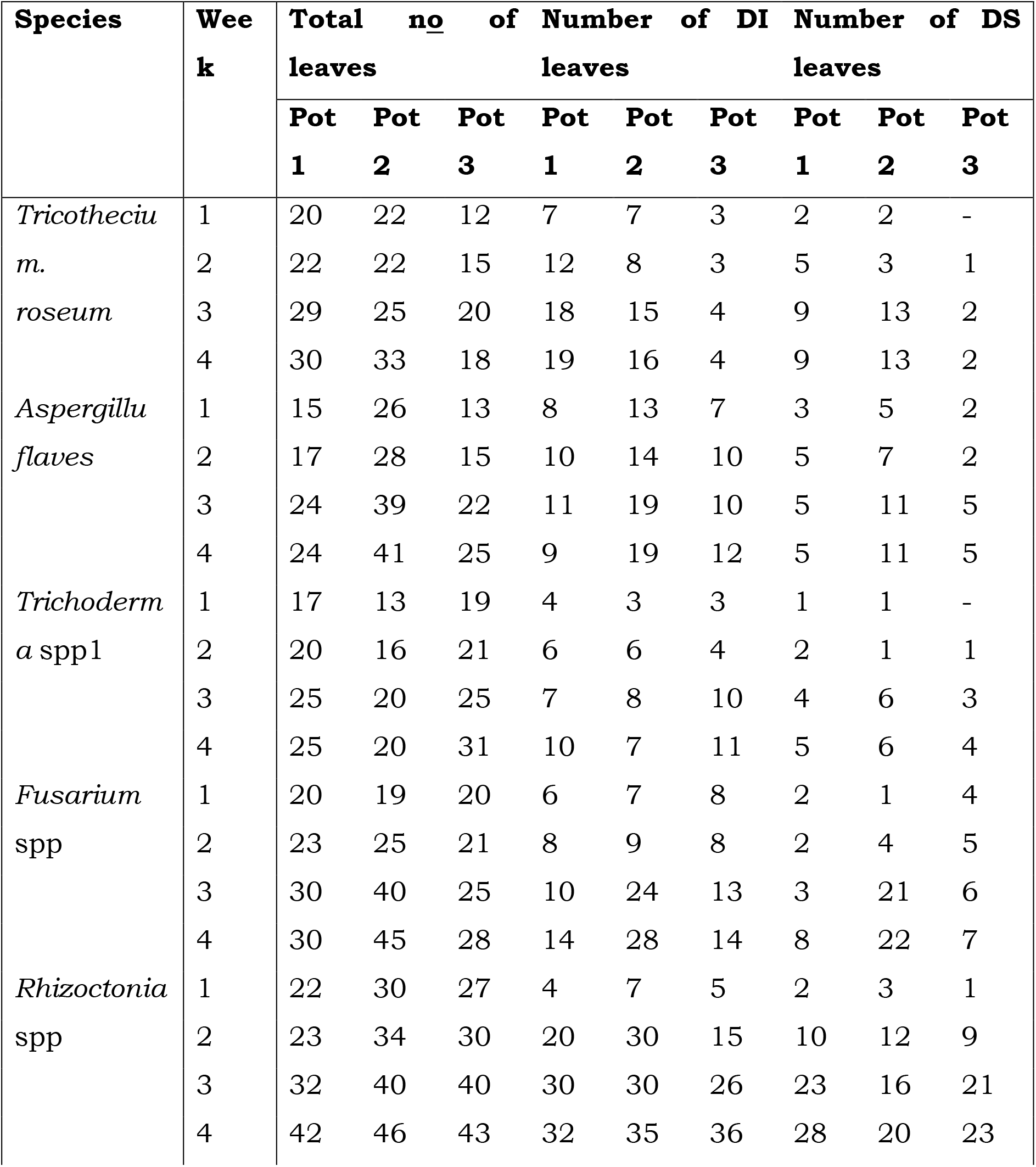
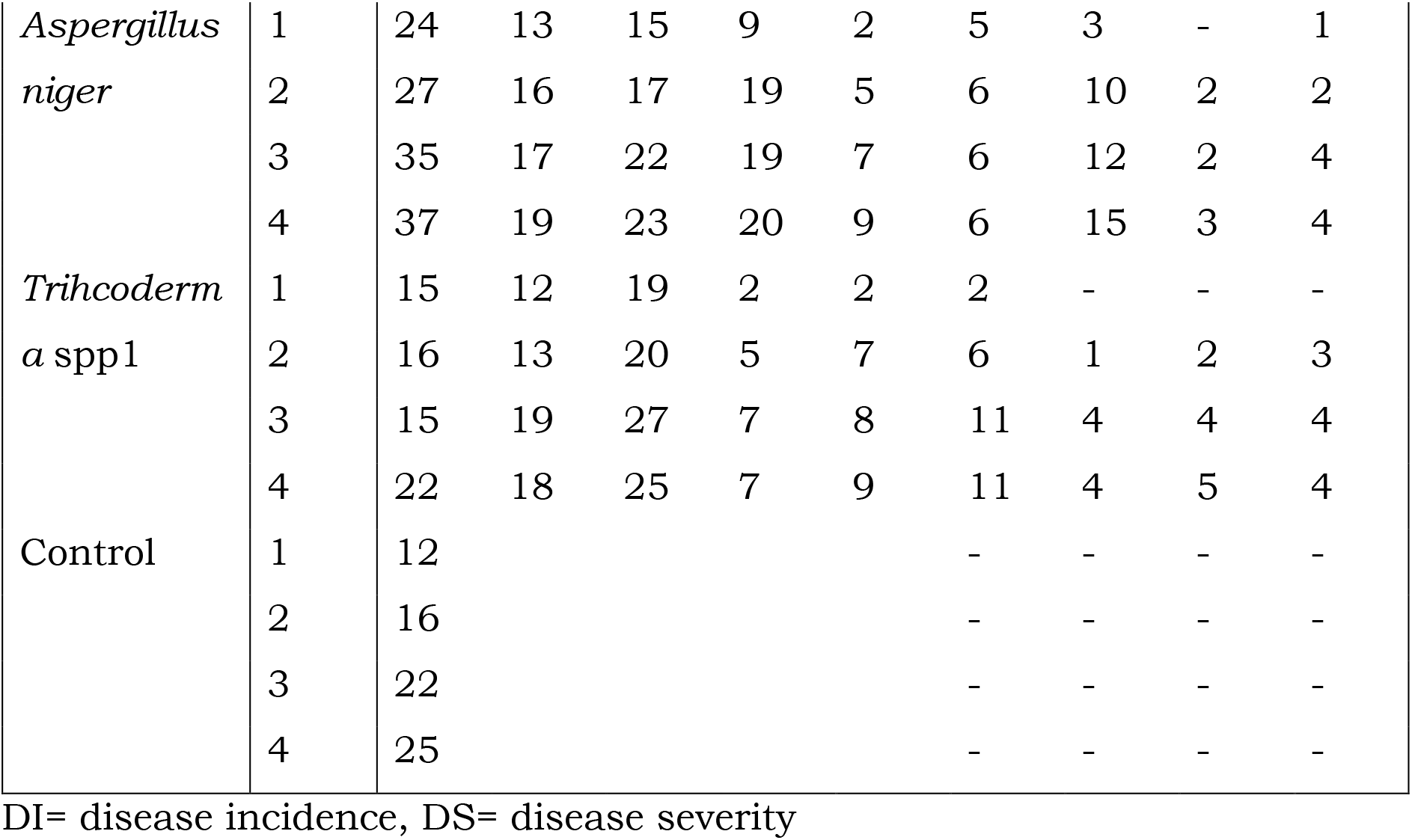
Level of Disease intensity effects of fungal isolates on water hyacinth under pot experiment conditions.

### 4.1. Disease incidence

At the end of the five weeks the highest disease incidence per plant was recorded in plants inoculated with *Rhizoctonia* spp (66.0 %). This was significantly higher than that of the *Tricothecumroseium* (50.7 %) (p=0.05) which was rated second followed by *Aspergilluflaves* (49.1 %) *Fusarium* spp (45.3 %) and *Aspergillus niger* (42.6 %). The least disease incidence was recorded in *Trichoderma* spp1 (34.2 %) and *Trichoderma* spp2 (31.3 %) significantly lower than the others. Disease symptoms cannot be observed on the control group.

### 4.2. Disease severity

On the basis of disease severity (DS) the pathogenic species were divided into less than 25 % mild, low moderate isolates caused 26 - 50 % damaged of the leaf area, high moderate isolates that damaged 51-75 % of the leaf area and severely fungi that cause greater than 75 % damage of the leaf area. Of these species only *Rhizoctonia* spp (100 %) and *Fusarium* spp (78.6 %) showed high disease severity. These fungi were associated with a high percentage of tissue death after five weeks application. *Aspergillus flaves, Tricothciumroseum* and *Aspergillus niger* show high moderate disease severity (58.3 %, 56.4 % and 53.6 %) tissue death, respectively. *Trichoderma* spp2 (31.4.7 %) and *Trichoderma* spp1 (27.7 %) shows low moderate. Most isolates showed minimum lesion growth during the first week. Control plants not show disease severity.

Also, the fungus that show a promising result in temporary shade and it have attempted in the open ponds that holds propagted healthy water hyacinth plants in September 2016 and the released fungus started attacked the water hyacinth when the season becomes hot. During rain and cold condition they become latent but,when the seasons became hot it attacked the water hyacinth on the ponds since October 2016 - January 2017.

**Figure 1:**
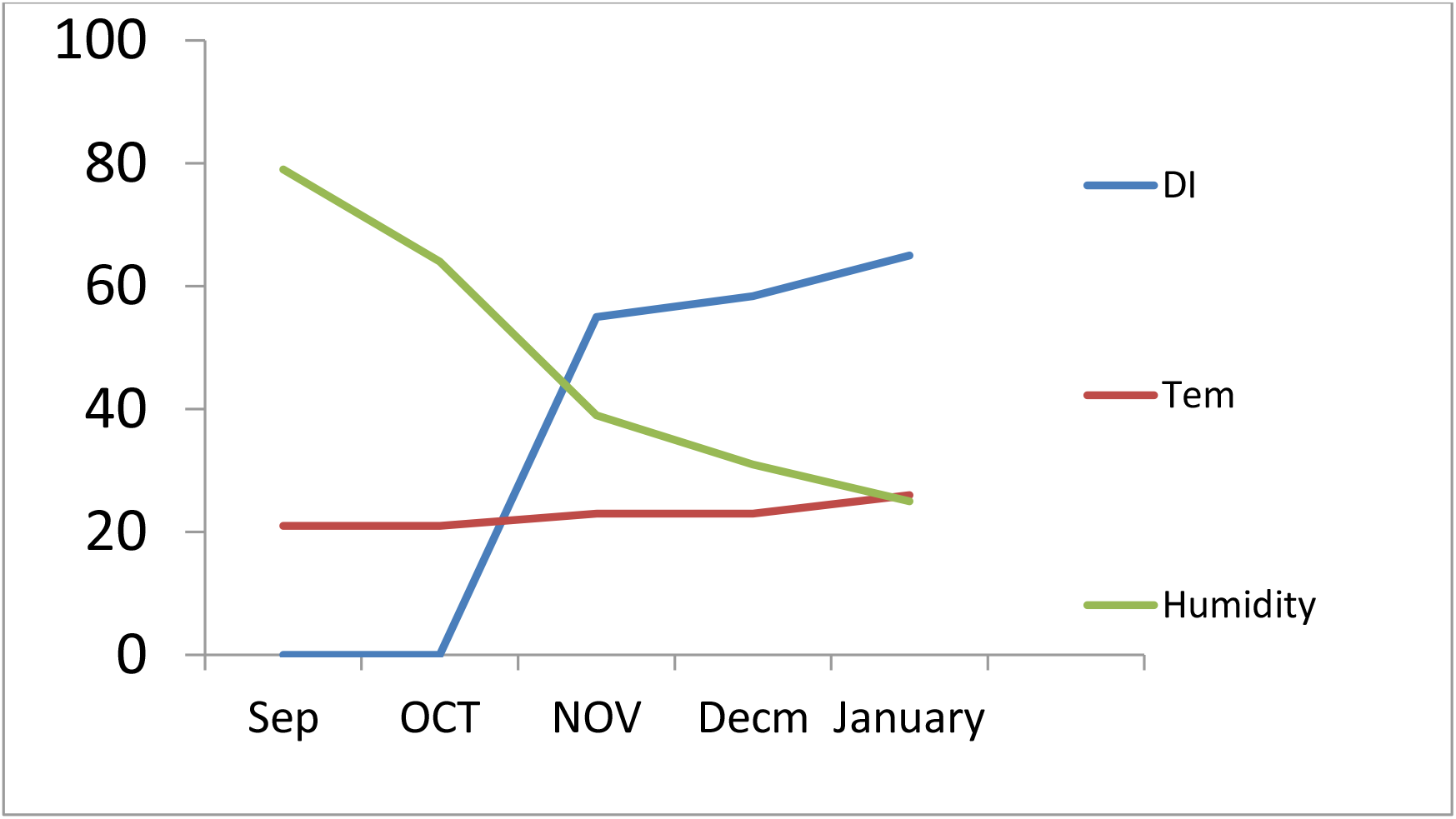
The fungus, Humidity and Disease severity association on water hyacinth lea**f** **Note:-**DI = disease insidence, Tem = Temperatur

**Table 2:** Modified Naseema et al., (2001) disease severity rating scale in 2016.

| R<br>S | Type of symptom produced/ symptom description | Nov | Dec | Jan | Feb |
| --- | --- | --- | --- | --- | --- |
| 1 | No symptom |  |  |  |  |
| 2 | 1-9 % symptom developed around the pin pricked area only |  |  |  |  |
| 3 | 10 % of the leaf area showing yellowing or browning |  |  |  |  |
| 4 | 11-25 % of the leaf area showing yellowing or browning |  |  |  |  |
| 5 | 26-50 % of the leaf area including petiole showing symptom | 41.6 % |  |  |  |
| 6 | 51-100 % of the leaf area including petiole showing symptom to complete drying of the plant |  | 52.4 % | 76.5 % | 100% |

Similarly, pathogencity tests of indigenous fungal pathogens on water hyacinth were tried in different countries. For instance, in Lake Victoria, Lake Naivasha and Nairobi Dam in Kenya were tried with 20 strains of pathogenic fungi. The Pathogenicity tests indicated that *Cercospora, Fusarium* and *Alternaria* spp. were diagnosed as potential mycoherbicides (Mailu et al., 1998) on water hyacinth. Martinezand Charudattan (1998) reported that *Alternaria* spp. and *Fusarium* spp. were highly virulent and severely damaged the inoculated water hyacinth leaves. Accordingly our research finding also related with this author because in our case *Fusariume spp*. was one of the promising biological control during temporary shade and pond trial.

**Figure 2.**
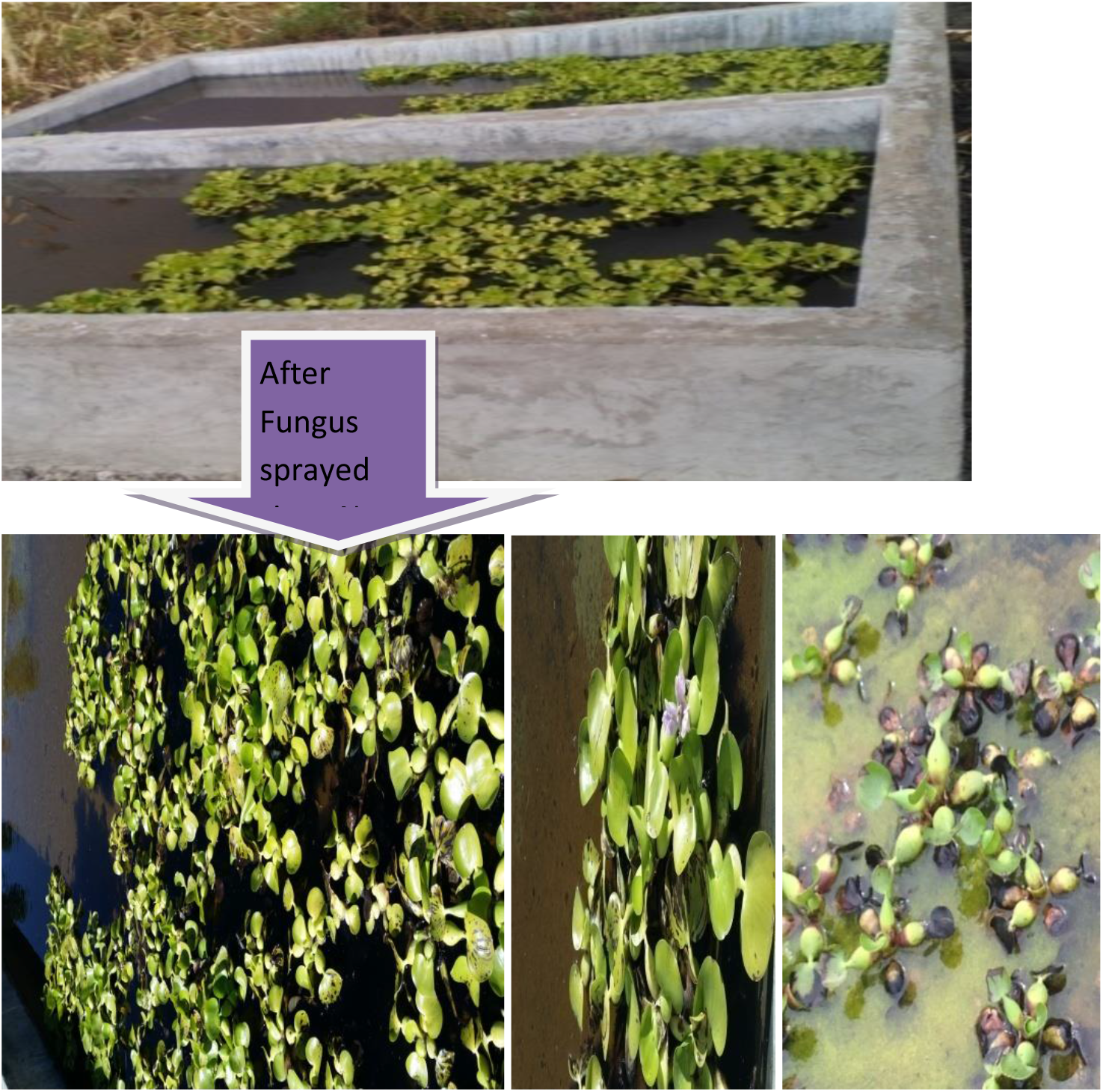
Treated by fungus on Feb.2017.

### 4.3. Study the impacts of *Rhizoctonia solani., Aspergillus flavus, Tricothcium roseum, Aspergillus niger on selective Lake Tana aquatic plants and fish*

Lake Tana basin is one of the three sections of Eastern Afromontane biodiversity hotspot (Mittermeier et al., 2004). It contains 65 fish species with about 70 % of the fish species in the lake being endemic (Goshu et al., 2010). Therefore, before directelly released to the Lake Tana the above mentioned biocontrols, this research study was assese its impact on most common biodiversirity Lake Tana by preparing 6×6 metre width and 1 metre depth of 3 open ponds.In those plot it was attempted to see the consequence of the fungus on some common biodivesity including Echinochloa and Cyperus papyrus, algae and barbus fishe of the Lake. By releasing those species togather with the biocontrols in the prepared ponds, for a year the impacts of the fungi was studied in this prepared site. But, there is no scientific data recorded that shows the impacts on those biodiversity. Finally, to investigate the effeciancy of the above mentioned biocontrols in wet land areas that are not linked with the lake were expermented by spraying 2ooo ml cultivated all mixed fungus in 20 metre square areas covered by water hyacinth in Goregora side of Lake Tana and it was effective.

**Figure 4.**
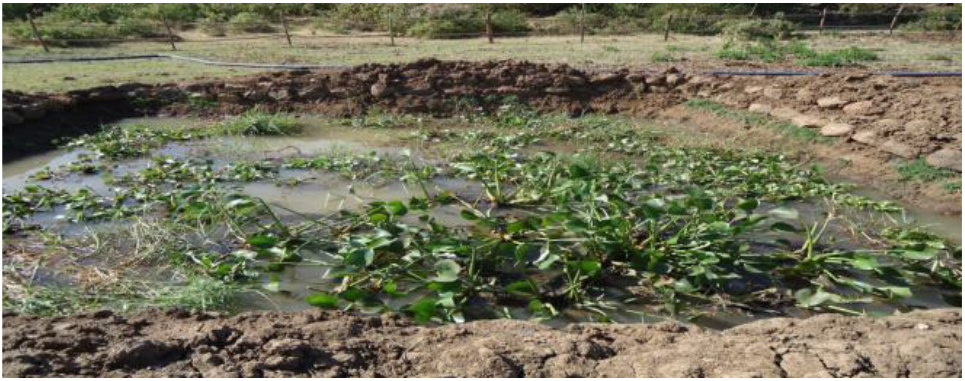
The impact studied pond on Dec.2017.

**Figure 5.**
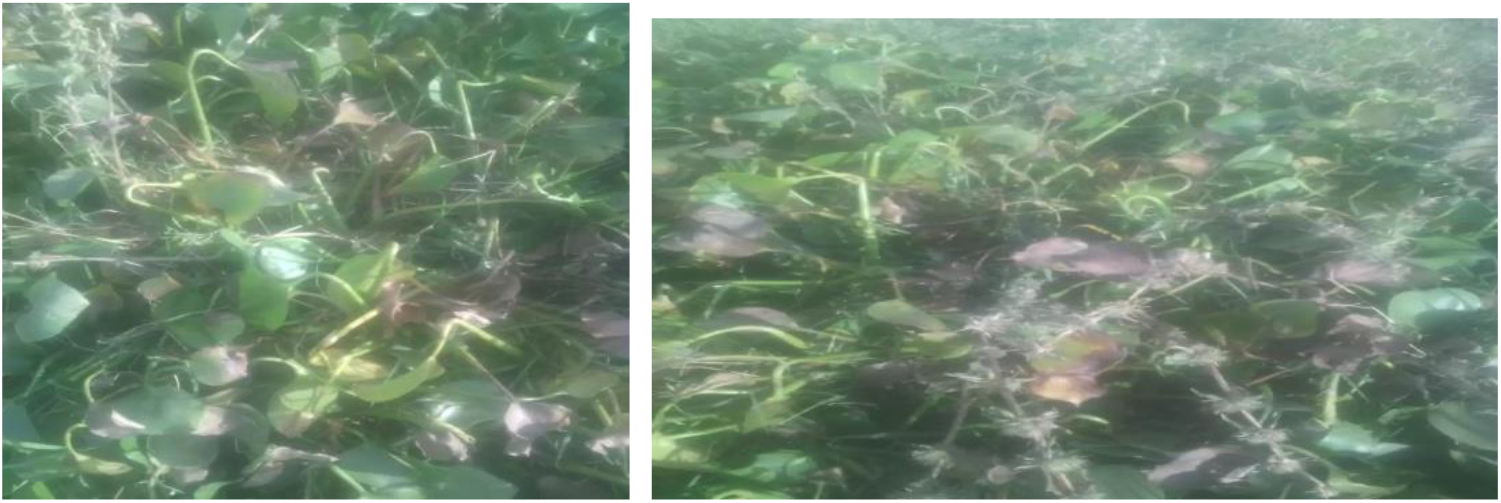
Treated water hyacinth by fungi in wetland of Lake Tana Nov. 2019.

**Figure 6.**
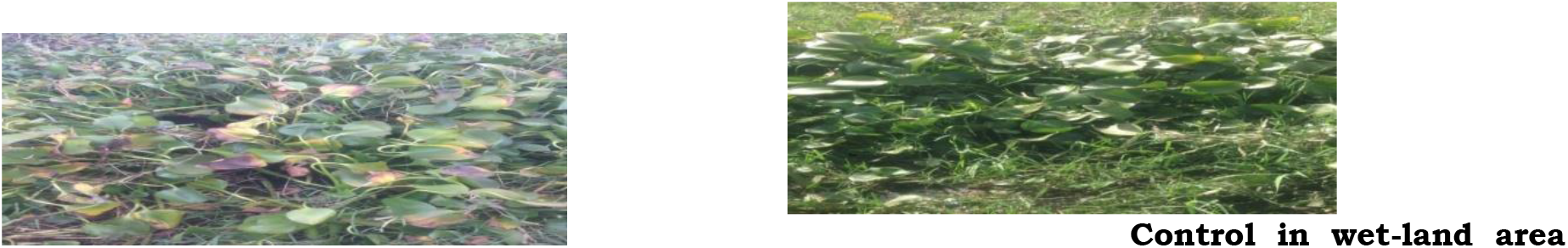
Treated water hyacinth by fungi in wetland of Lake Tana Dec 2019.

## 5. CONCLUSION AND RECOMMENDATION

Tricothecium roseum, Aspergillus Flaves, Trichoderma spp1-,Fusarium spp, Rhizocotonia spp, Aspergillus niger and trichdoerma spp fungi were synregetically promising to minimize water hyacinth expansion at above 26 °C and at less than 25% humidity.Also, those fungi have not any impact on aquatic biodiversity of the Lake Tana. Because, most of the lake is thretened by water hyacinth in Dembeya, Maksegnete and Fogera dstrictes since this places of the Lake edge have accessed for erosion soils because this erosion hold N and P nutrient.This nutrients make a good opportunity for spreads of water hyacinth, but the lake that surrounds by Bahrdar city have not observed this weed because it has not get the chance to get favorable nuteriant source, so, if urbanization is expaneded in all Tana lake edges the lake will protected from any enviroment pollutanet since at the time internationally standardized buffer zone is established and the lake will never have a chance to connect with any enviroment pollutanet source.

## 6. ACKNOWLEDGEMENTS

Ethiopian Environment and forest research institute, DebereTabor University and University Gondar would like to be acknowledged for their financial support to conduct the research.

